# Report of unexpected findings after cardiac stem cell injections in a preclinical model

**DOI:** 10.1101/2021.10.26.465939

**Authors:** Mira van der Naald, Hans T van den Broek, John LM Bemelmans, Klaus Neef, Maarten H Bakker, Patricia YW Dankers, Adriaan O Kraaijeveld, Steven AJ Chamuleau

## Abstract

**Introduction:** Cardiac regenerative therapy is a proposed therapy for ischemic heart disease. So far efficacy has been low and this might partly be explained by low cardiac cell retention. In this study we aimed to investigate if cardiac cell retention improves using ureido-pyrimidinone units (UPy-gel) as a cell carrier.

**Methods:** We used an ischemia-reperfusion model. Pigs were randomized to intramyocardial injections with mesenchymal stromal cells (MSC) labelled with both Indium-111 and a fluorescent tracer in either PBS or in the UPy-gel. After 4 hours, a total body scintigraphy was performed to determine the cardiac cell retention and histology was obtained.

**Results:** In the first 4 pigs, we noticed focused areas of radio activity (hotspots) outside the heart in both the control and UPy-gel arm, and decided to interrupt the study. At histology we confirmed one hotspots to be located in a lymph node. No satisfactory explanation for these, potentially harmful, hotspots was found.

**Conclusion:** This study was interrupted due to unexpected extra-cardiac hotspots. Although we do not have a conclusive explanation for these findings, we find that sharing these results is important for future research. We recommend to use total body imaging in future retention studies to confirm of reject the occurrence of extra-cardiac cell accumulation after intramyocardial cell injection and discover the pathophysiology and its clinical implications.

## INTRODUCTION

Cardiac cell therapy has been a promising therapy to repair the damaged heart. However, efficacy has been low in preclinical and clinical trials^1,2^. One possible explanation for the observed low efficacy could be inefficient cell delivery. We previously showed that cardiac retention after intracoronary infusion or intramyocardial injection of bone marrow derived mesenchymal stromal cells (MSC) is limited to 10-15%^3,4^. Additionally, we showed that retrograde coronary venous infusion does not improve cardiac retention^4^. In this study we aim to test if delivery with a cell carrier improves cardiac retention. Here we use a pH-switchable hydrogel based on ureido-pyrimidinone units telechelically coupled to poly(ethylene glycol) (UPy-gel)^5^. This hydrogelator is in the liquid state at basic conditions and turns into a gel state at a lower, i.e. neutral or acidic, pH. We aimed to show increased cardiac retention when injecting MSCs combined with UPy-gel, compared to MSCs in phosphate-buffered saline (PBS) in a confirmatory pig study. We found extra-cardiac focused areas of high intensity signal (hotspots) implying extra-cardiac accumulation of cells in the first pig and confirmed this in the following 3 pigs. The hotspots were observed in both study arms. This finding was unexpected and has potential harmful clinical consequences. Therefore we decided to interrupt and de-blind this study. Here we share our unexpected findings, discuss possible explanations and provide recommendations for future research.

## METHODS

### Ethical statement

All experiments were performed in compliance with the “*Guide for the Care and Use of Laboratory Animals*”, published by the National Institutes of Health (National Institutes of Health publication 85-23, revised 1985). The protocol was approved by the Animal Experiments Committee of the Utrecht University (AVD115002015257) and registered at www.preclinicaltrials.eu (PCTE0000105). Protocols of comparable experiments are available online^3,4,6,7^.

### Animals and housing

Female Yorkshire pigs (van Beek, SPF varkensfokkerij B.V. Lelystad) of approximately 70 kg were used in these experiments. Animals were housed in stables embedded with straw and enriched with rods. Animal welfare was assessed on a daily base by animal caretakers.

### Study design

Myocardial infarction was induced at baseline. After 4 weeks, all surviving pigs were randomized to intramyocardial injections of mesenchymal stromal cells (MSC), radioactively labeled with Indium^111^ and fluorescently-labeled with carboxyfluorescein succinimidyl ester (CSFE), in either a solution of 1) PBS or in 2) UPy-gel (figure 1). If animals reached an human endpoint (severe immobility, severe dyspnea or cyanosis, wound infection) they were euthanized and excluded. There were no additional inclusion criteria. According to sample size calculations, 14 pigs were needed to show a 6% increase in cardiac cell retention. The alpha was set on 0.05, beta on 0.20, the standard deviation on 3 and we expected 20% of the animal to drop-out due to fatal rhythm disorders during or shortly after infarct induction. We used block randomization, generated by a computer-generated random number sequence. Animals were randomized in a one-to-one ratio. All procedures were performed by the same researchers (cell culture (KN), catheter handling (MN), cell labeling and syringe control (TB)). The researcher handling the catheter was blinded for treatment allocation. Scintigraphy analyses, including drawing the regions of interest in the scintigraphy images, were performed by the same two technicians and supervised by the same nuclear medicine physician (JB), all of them were blinded for treatment allocation.

**Figure 1:**
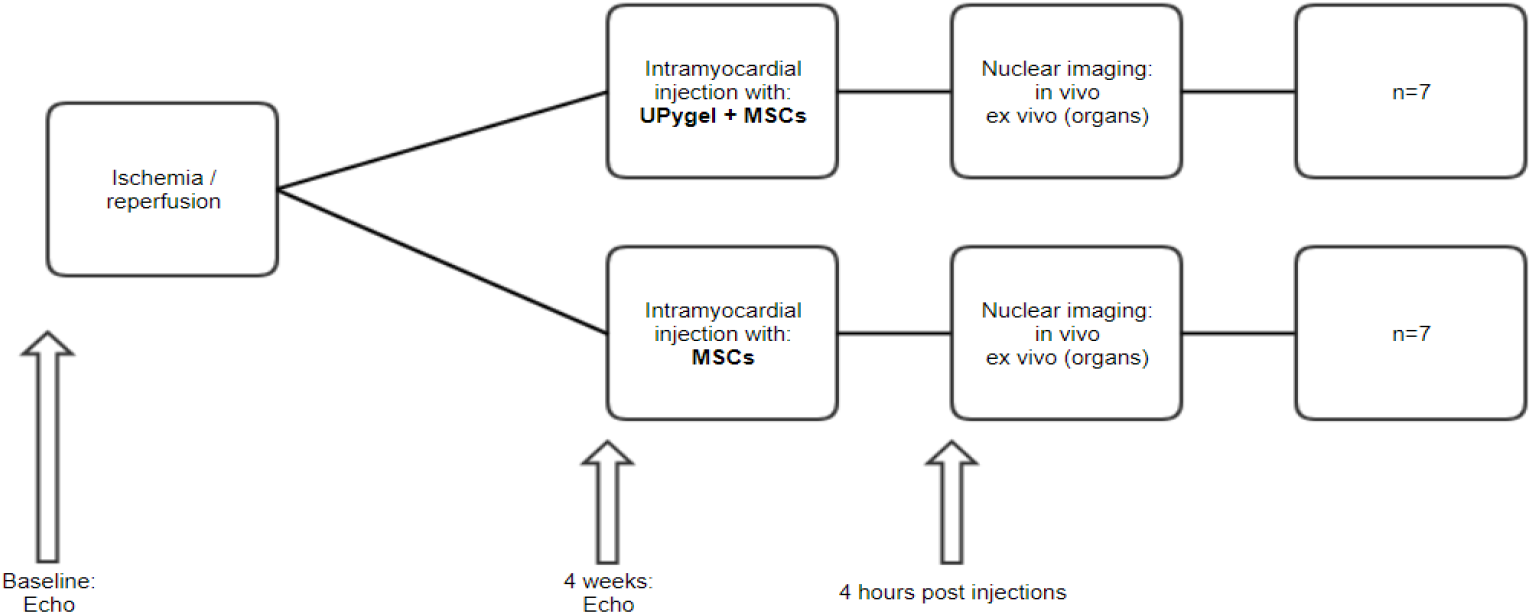
Study design. Ischemia/reperfusion was induced by a 90 minute occlusion of the Left Anterior Descending artery with a balloon via a percutaneous procedure. Four weeks after ischemia-reperfusion, intramyocardial injections were performed. Four hours after injections in vivo total body scintigraphy was performed, and the pigs were sacrificed for ex vivo scintigraphy of the organs and histology.

### Anesthesia and analgesia

All animals were treated with amiodarone (1200 mg/day, 7 days), clopidogrel (75 mg, 3 days) and carbasalate calcium (320 mg, 1 day) prior to the myocardial infarction. Animals were anesthetized in the supine position with intramuscular ketamine (10-15 mg/kg), midazolam (0.7 mg/kg) and atropine (0.5 mg) and intravenous thiopental sodium (4 mg/kg), midazolam (10 mg) and sufentanil (0.25 mg). A bolus of amiodarone (300 mg in 30 minutes) was administered intravenously. During the procedure the animals received midazolam (1mg/kg/h), sufentanil 10 µg/kg/h) and pancuronium bromide (0.1 mg/kg/h). Heparin (5000 IU) was given every 2 hours. All animals received a butrans patch (5 µg/h). Animals were ventilated with a mixture of dioxygen (O2) and air (1:2) with a tidal volume of 10 ml/kg with 12 breaths per minute. Carbasalate calcium was continued (80 mg/day) until euthanasia.

### Ischemia-reperfusion model

Animals were monitored during the entire procedure via continuous electrocardiogram, arterial pressure and capnogram. First the left coronary system was visualized via a coronary angiography. The myocardial infarction (MI) was induced by a 90-minute occlusion of the left anterior descending artery (LAD) using an angioplasty balloon. The balloon position was based on the coronary anatomy, the preferred position was after the second diagonal branch. In case of ventricular fibrillation or ventricular tachycardia without output, an electrical shock of 200 joules was delivered using an external defibrillator. Additionally, chest compressions were given and animals received amiodarone (150 mg, max 3 times), adrenaline (0.1 mg) and/or atropine (0.5) mg.

### Cell culture and labeling

For this experiment we used allogeneic mesenchymal stromal cells (MSCs). These were isolated from the sternum and cultured as described earlier^8^. Cells (1 × 10^7^) from passage 5-7 were used for transplantation after staining with carboxyfluorescein succinimidyl ester (CSFE) (Invitrogen, Carlsbad, California, USA) on the day of the transplantation. Cells were labelled with 30 MBq In^111^ by incubation at 37°C for 20 minutes and washed with Hank’s balanced salt solution (Life Technologies Corp, Grand Island, New York, USA) to remove excess unbound In^111^ as described before^3^.

### Hydrogel specifications

The UPy-hydrogelater (SyMO-Chem BV, Eindhoven, the Netherlands) was prepared as described before^5,9,10^. In short, the UPy-hydrogelator was dissolved at 5 weight percentage (wt%) in phosphate buffered saline (PBS) pH 11.7 and temperature of 70 °C using a magnetic stirrer. After dissolving, the solution reaches a pH of 9.5. The solution was then cooled down. The cells were then pipetted into the solution and stirred for 10 minutes to reach uniform distribution.

### Intramyocardial cell injection

An electromechanical map of the left ventricle was obtained using the NOGA system (Biosense Webster, Cordis, Johnson & Johnson, USA). Cells were injected in the myocardial border zone as previously defined, using the MYOSTAR® injection catheter (Biosense Webster, Cordis, Johnson & Johnson, USA)^11^. Per injection approximately 0.3 mL was injected, 10-12 injections were performed per pig. Needle depth was set at 5-7 mm. The cells were injected slowly, approximately 30 seconds per injection, and the injection needle was left in situ for an additional 10 seconds to avoid leakage.

### Nuclear imaging and analysis

A scintigraphy scan, using a dual head gamma camera (Philips NM SkyLight) was performed after 4 hours to determine cell retention in the heart and other organs of interest (liver, spleen, kidneys, lung, and bladder) (figure 1). First, an in vivo total body scan was performed at 174 keV and 247 keV energy windows. After euthanizing the animal, the organs of interest were excised and scanned. Anterior and posterior images were captured for the total body scan and the ex-vivo scan of the organs. The number of counts was based on the geometrical mean of the anterior and posterior counts. Cell retention was measured by the number of counts in the region of interest as a percentage of total body activity. Analysis were performed directly after each experiments by a team blinded to treatment allocation.

## RESULTS

We performed experiments with 4 out of 14 pigs according to protocol, with an experienced team and did not encounter any obvious technical issues. After analyses of our first results we found focused areas of radio-activity (hotspots) outside the heart (figure 2). These hotspots were distributed throughout the body, including the abdomen, head and extremities. We did not expect to find any hotspots outside the target organs, and suggested this can compromise the value of this study. We decided to interrupt and de-blind the study after 4 pigs to investigate a reasonable explanation for the origin of these hotspots. Since we could not find a satisfying explanation and could not rule out potential harm of these hotspots, we decided to stop the study. Ethical considerations regarding use of animal and resources also contributed to this decision.

**Figure 2:**
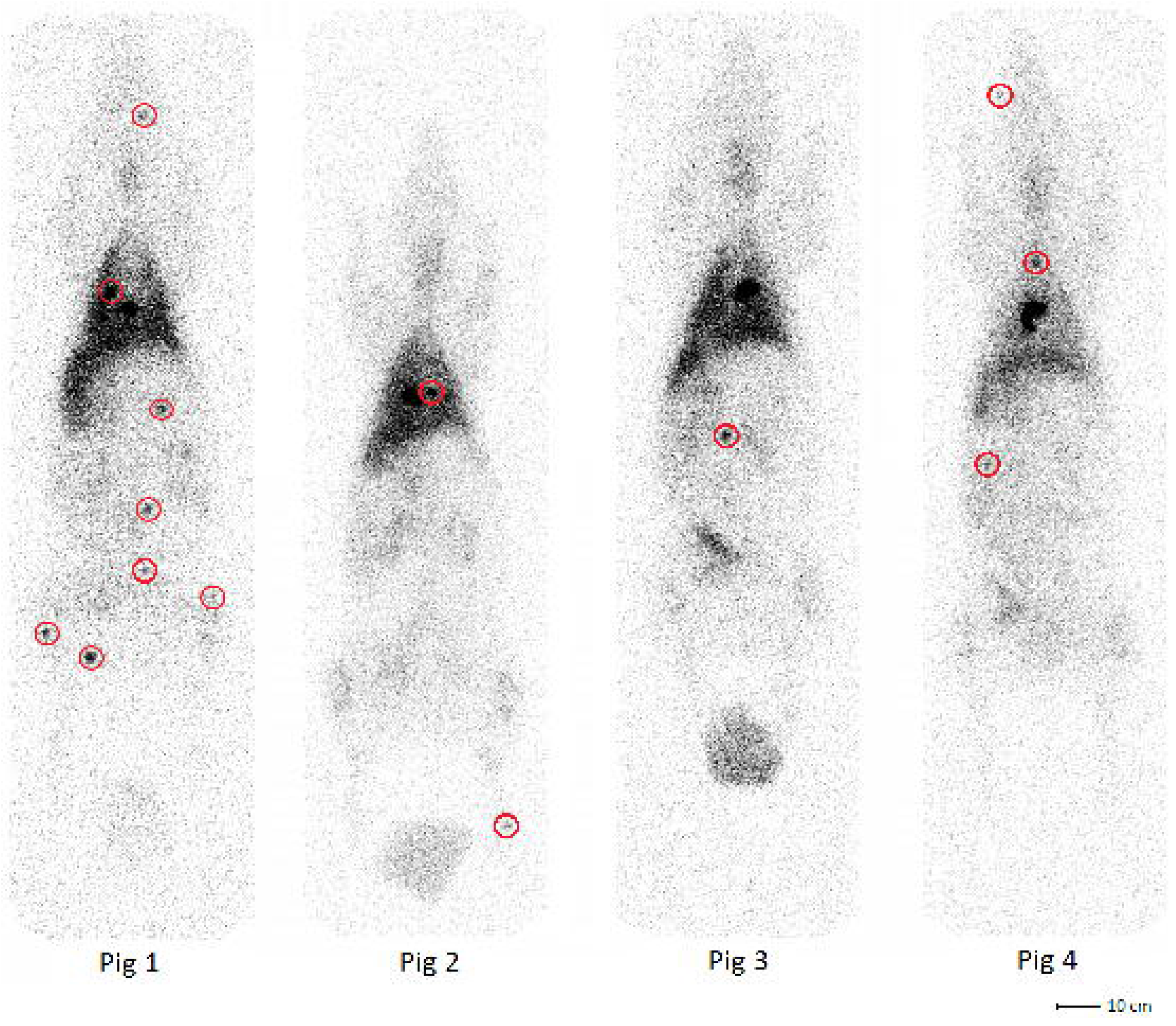
Total body (including urine catheter) scintigraphy scan images 4 hours after injection. Pig 1 and pig 4 were randomized to UPy-gel injections, pig 2 and pig 3 were injected with cells in PBS. The hotspots are marked with red circles.

### Hotspots

Two authors (TB and MN) discussed the scintigraphy images and rated areas of increased signal intensity as hotspots by visual inspection. Quantification of signal intensity over background in the hotspots did not occur. In the UPy-gel group we identified a total of 11 hotspots (8 and 3), compared to 3 hotspots in the PBS group (2 and 1). We tried identifying the exact location of the hotspots by obduction and with use of the scintigraphy scan. We traced one of the hotspots to a lymph node. However not all hotspots were traceable with this strategy. Histology confirmed CSFE-labelled MSCs in the retrieved hotpot (figure 3). Unfortunately, we could not perform additional imaging (i.e. computed tomography scan) within this study.

**Figure 3:**
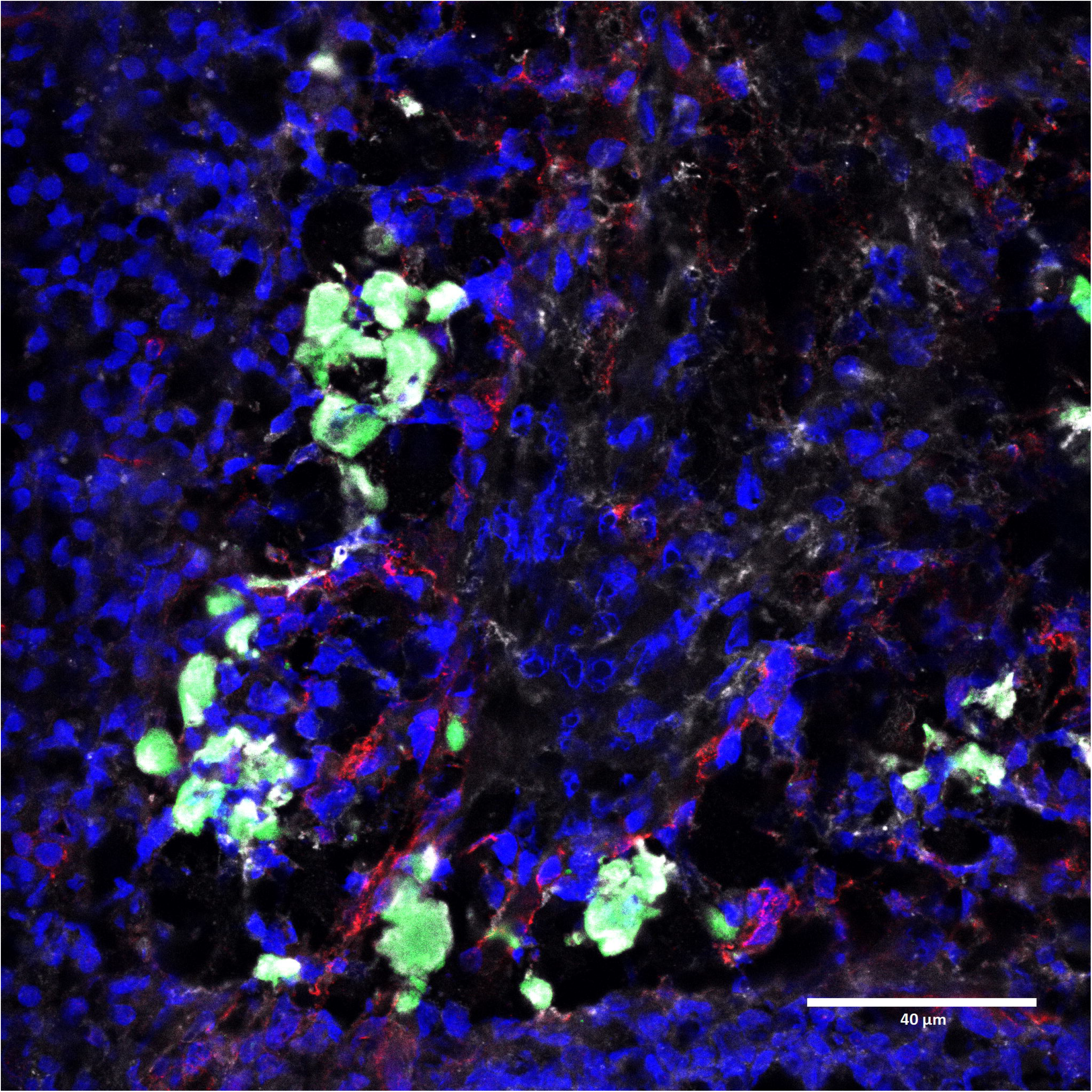
Histology performed on one hotspot (lymph node): Green: CSFE labeled injected MSCs. Red: CD31 endothelial vascular cells. Blue: Hoechst nuclei. Gray: Ly6G immune cells.

### Cardiac retention

Whole body scintigraphy revealed that cardiac retention was low in both groups. Retention in the heart was 4.3% and 5.3% in the UPy-gel group compared to 3.4% and 4.0% in the PBS group (table 1). Cells accumulated in lungs, liver, kidney and spleen.

**Table 1:** Cell retention in the target organs, measured as number of counts as percentage of number counts in the total body

|  | Heart | Lungs | Kidneys | Liver | Spleen |
| --- | --- | --- | --- | --- | --- |
| Pig 1 (UPy) | 4.3% | 17.2% | 2.7% | 8.2% | 1.6% |
| Pig 2 (PBS) | 3.4% | 18.8% | 3.2% | 9.5% | 0.7% |
| Pig 3 (PBS) | 4.0% | 23.1% | 2.9% | 4.2% | 1.1% |
| Pig 4 (UPy) | 5.3% | 20.4% | 2.8% | 4.2% | 1.0% |

### Tracing of UPy-gel

We hypothesized that the UPy-gel would turn into a gel state immediately after injection and thus remain in the heart as previously shown^5,10,12^. We further hypothesized that the UPy-gel might have remained in the heart and only the radio-active labeled cells were distributed throughout these hotspots. We therefore performed an additional, post-hoc, in vivo experiment (n=1) to investigate whether hotspots contain UPy-gel. UPy-gel (5 wt%, pH 9.5) in combination with UPy-DOTA-Gadolinium (UPy-DOTA), which is traceable with magnetic resonance imaging (MRI), was injected in combination with radioactive labeled MSCs via intramyocardial injections, using the same number of cells and injection method as the original experiment^12^. Scintigraphy showed 4 intra-cardiac hotspots and 1 extra-cardiac hotspot in the mediastinum (figure 4A). An MRI of the heart confirmed the intra-cardiac hotspots contained UPy-gel. No additional imaging techniques or imaging of the extra-cardiac hotspot were performed in this experiment.

**Figure 4A (left):**
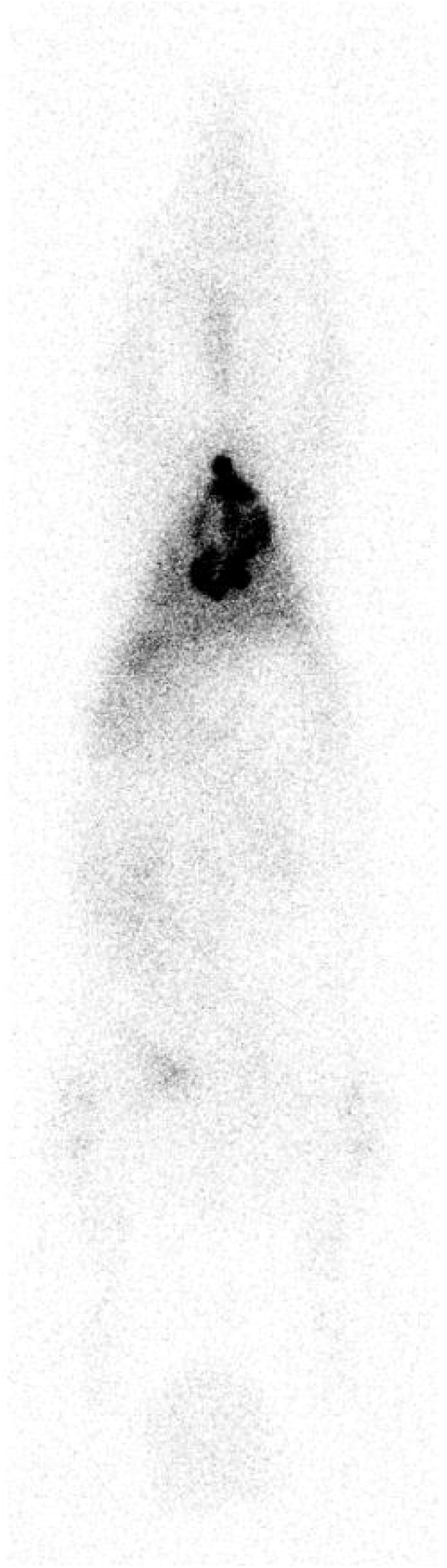
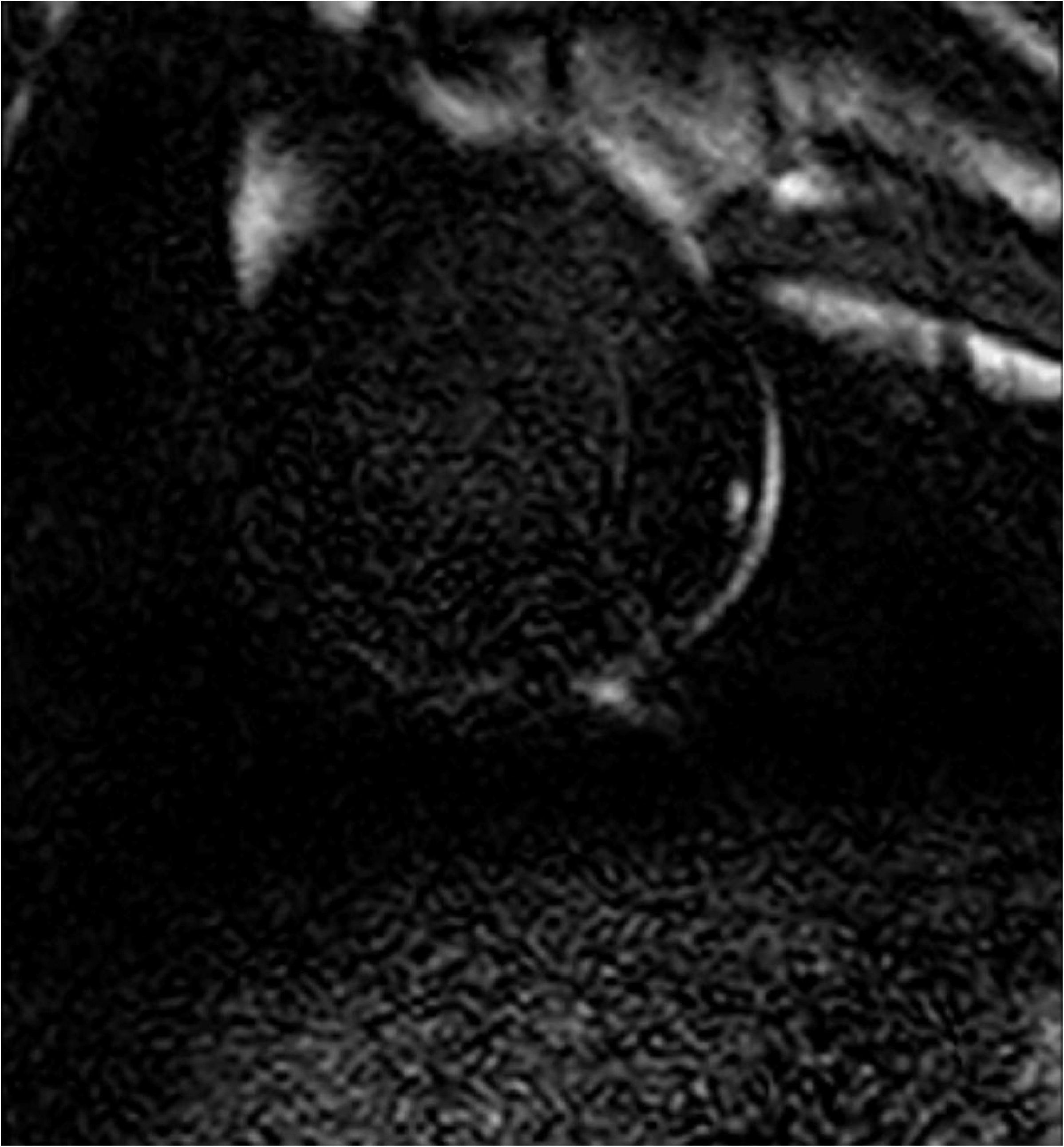
Scintigraphy image of post-hoc experiment with 1 pig using UPy-DOTA. Figure 4B (right): Short axis 3D viability scan with SENSE of post-hoc experiment with 1 pig using UPy-DOTA.

## DISCUSSION

With this study we aimed to show increased cardiac retention of cells using a cell carrier in an animal model. We found extra-cardiac hotspots in the first 4 out of 14 pigs, in both the PBS and the UPy-gel group. Additionally, the cardiac retention in these four pigs was lower than expected based on previous experiments using the same protocols. We could not find a satisfactory explanation for these findings and propose these results potentially compromise the value of this study. Therefore we decided to interrupt this study. Here we share our unexpected findings, not only because we find sharing (unexpected) results contributes to transparent research, but we also propose these findings demand further research to confirm the safety of intramyocardial cell injections in this model.

### Extra-cardiac hotspots

Tracing of cells after cardiac transplantation has been performed in several animal studies and a little number of clinical studies. Based on these previous studies, we know that cardiac retention is low and most cells can be traced back in the lungs, intestine, kidney, bladder and liver ^3,13–16^. We expected to find diffusely distributed radio-activity outside the heart. Surprisingly, in the present study we found focused areas of radio-activity outside the heart (hotspots). Four potential explanations were considered: arterial embolisms, role of the hydrogel, venous-lymphatic spill, or technical issues. First, the cells could have formed clots in the myocardium and leak back in the left ventricle (or pushed out of the myocardium by cardiac contraction) through the injection site, causing potential harmful arterial embolisms. We could not rule out arterial obstructions in this study as we did not perform CT-angiography. Importantly, in clinical studies over 2600 people received cardiac cell transplantation, of which over 200 patients received percutaneous intramyocardial cell injections. In these studies cell therapy seems to be safe and did not show a major risk of embolisms^17^. Second, we considered the hydrogel to contribute to these hotspots. We found hotspots in the study arm without the use of this hydrogel. We re-analyzed data of our previous retention study with intramyocardial injections of mesenchymal stromal cells in PBS with a comparable study protocol, but without the use of a hydrogel carrier^3^. Although this was not reported specifically, in hindsight hotspots were also visible. Taken together, we propose that is it is unlikely that the hydrogel plays a role in the formation of hotspots. Third, we hypothesized that the cells could have entered the venous system of the heart. Involvement of the lymphatic system is suggested to explain the prominently right-sided distribution of cells^18^. Possibly, the lymphatic system could then play a role in formation of hotspots, as we confirmed one extracardiac hotspot to be located in a lymph node. A clinical study that traced cells and performed total body imaging after intracoronary infusions, which is expected to have comparable venous drainage, did not show any extra-cardiac hotspots and could not provide evidence of involvement of the lymphatic system^15^. The fourth explanation could be technical issues. We have a team of skilled technicians and researchers with abundant expertise in translational studies for cardiac regeneration. Experiments are conducted according to strict protocols^6,7^. With these measures we limited the risk of a procedural flaw. Hotspots were, when looking back at previous work, only found in studies with intramyocardial injections. We considered the possibility of a technical failure of these injection catheters. High pressure is used to inject the product through the catheters, that potentially could have led to failure (e.g. damaged lumen or damaged injection needle). However, we exclude such technical issue since we checked and flushed all catheters after the procedures and did not find any problem/inconsistency.

Three additional studies were found that performed percutaneous intramyocardial cell injections and performed total body imaging (table 2)^13,14,19^. All three studies were performed in pigs and used the same MYOSTAR® catheter to perform cell injections. Collantes et al applied positron emission tomography/computed tomography (PET-CT), allowing 3D visualisation of all tissues^14^. This study describes high radioactivity concentrations in mediastinal lymph nodes. Perin et al used a reporter gene, which passes on to daughter cells during proliferation, and performed repetitive imaging over time. They described involvement of the lymphatic system around the heart and cervical region ^19^. It should be noted that the distribution of the hotspots seems to be different in our study, as not all hotspots in our study are located in the mediastinum. Nevertheless, this supports one of our theories that the lymphatic system plays a role. Interestingly, Lyngbæk et al did not report extra-cardiac hotspots^13^. A CT-angiography to rule out arterial embolisms was not performed in any of these studies.

**Table 2:** Comparison of studies on in vivo cell tracking, all studies are performed in pig models. I/R = ischemie/reperfusion, PET-CT= positron emission tomography-computed tomography.

|  | Present study | van der Spoel<sup>3</sup> | Lyngbæk<sup>13</sup> | Collantes<sup>14</sup> | Perin<sup>19</sup> |
| --- | --- | --- | --- | --- | --- |
| **Porcine model** | I/R | I/R | Healthy | I/R | I/R |
| **Cells** | Mesenchymal stromal cells | Mesenchymal stromal cells | Mesenchymal stromal cells | Cardiac stem/progenitor cells | Mesenchymal stromal cells |
| **Cell donor** | Allogeneic | Allogeneic | Xenogeneic (human) | Allogeneic | Autologous |
| **Number of cells** | 1 x 10<sup>7</sup> | 1 x 10<sup>7</sup> | 1.5 to 3.3 x 10<sup>6</sup> | 50 x 10<sup>6</sup> | 1 x 10<sup>8</sup> |
| **Label used** | Indium<sup>111</sup> | Indium<sup>111</sup> | Indium<sup>111</sup> | 18F-FDG/GFP | sr39HSV1-tk gene |
| **Volume injected** | 10-12 injections, 0.3 ml per injection | 10-12 injections, 0.3 ml per injection | 10 injections, 0.3 ml per injection | 30 injections, 0.3 ml per injection | 3 injections, 0.1 ml per injection |
| **Imaging technique** | Scintigraphy | Scintigraphy | Scintigraphy | PET-CT | [18F]FEAU PET/CT |
| **Timing of imaging** | 4 hours after injections | 4 hours after injections | 0.5 hour after injection | 4 hours after injections | 4 hours to 5 months after injection |
| **Hotspots outside target organs** | Yes | Yes | No | Yes | Yes |
| **Explanation for hotspots** | One in lymph node, other unconfirmed | No | Not applicable | Mediastinal lymph nodes | Periaortic lymphatic structures, coronary trunks, cervical lymph nodes. |

## Relatively lower cardiac retention

We observed in these 4 pigs that the cardiac retention is limited (3-5%), both in our control and UPy-gel group, and lower compared to previous work^3,4,13,14^. Clearly, this study was not completed and no definite conclusions can be drawn about cardiac retention. We did not find a clear explanation for the assumed lower cardiac retention. The risk of insufficient internal study validity (because previous results were not reproduced in our control group) contributed to the discussion to interrupt this study.

## Conclusion

This study was initially designed to show an increased cardiac retention with the use of a hydrogel, but was interrupted due to unexpected findings. We found extra-cardiac hotspots and a lower cardiac retention in our control group as expected. Although we do not have a conclusive explanation for these findings, we find that sharing these results are important for future research and contributes to transparency. Clinical trials did not show safety issues related to intramyocardial cell injections, but only a limited number of studies performed total body imaging and therefore extra-cardiac hotspots could have been missed. The limited number of studies that did perform total body imaging are all preclinical studies and have conflicting results. Most studies showed involvement of the lymphatic system, but the distribution of cell accumulation seems to differ from our current findings. Further research should confirm or exclude the occurrence of extra-cardiac hotspots after intramyocardial cell injection and provide a better understanding of its pathophysiology and clinical implications, before continuing research to optimize cell retention with carriers. We encourage researchers to include total body imaging in future research in this field.

## Acknowledgements

The authors thanks Marlijn Jansen, Joyce Visser, Martijn van Nieuwburg, Helma Avezaat and Jeroen van Ark for their excellent technical assistance and animal care. The authors also thank Monique Jacobs and Ingrid Boots for their work on the scintigraphy scans; Evelyn Velema and Joris de Brouwer for all planning and logistic affairs.

## Funding

This research is part of Cardiovasculair Onderzoek Nederland (grant number CVON 2011-2), an initiative of the Dutch Heart Foundation, Netherlands Federation of University Medical Centres (NFU), Royal Netherlands Academy of Arts and Science (KNAW) and NWO/ZonMW.

No human studies were carried out by the authors for this article.

All institutional and national guidelines for the care and use of laboratory animals were followed and approved by the appropriate institutional committees.

## ABBREVIATIONS

UPy-gel: ureido-pyrimidinone hydrogel
LAD: left anterior descending artery
MSC: mesenchymal stromal cells
PBS: phosphate-buffered saline
CSFE: carboxyflueroescin succinimidyl ester
MI: myocardial infarction
PET-CT: positron emission tomography/computed tomography
CT: computed tomography
FDG: fluorodeoxyglucose

## References

1. Jansen of Lorkeers SJ, Gho JMIH, Koudstaal S, et al. Xenotransplantation of Human Cardiomyocyte Progenitor Cells Does Not Improve Cardiac Function in a Porcine Model of Chronic Ischemic Heart Failure. Results from a Randomized, Blinded, Placebo Controlled Trial. PLoS One. 2015;10(12):e0143953. doi:10.1371/journal.pone.0143953

2. Gyöngyösi M, Wojakowski W, Lemarchand P, et al. Meta-Analysis of Cell-based CaRdiac stUdiEs (ACCRUE) in patients with acute myocardial infarction based on individual patient data. Circ Res. 2015;116(8):1346–1360. doi:10.1161/CIRCRESAHA.116.304346

3. van der Spoel TIG, Vrijsen KR, Koudstaal S, et al. Transendocardial cell injection is not superior to intracoronary infusion in a porcine model of ischaemic cardiomyopathy: a study on delivery efficiency. J Cell Mol Med. 2012;16(11):2768–2776. doi:10.1111/j.1582-4934.2012.01594.x

4. Gathier WA, van der Naald M, van Klarenbosch BR, et al. Lower retention after retrograde coronary venous infusion compared with intracoronary infusion of mesenchymal stromal cells in the infarcted porcine myocardium. BMJ Open Sci. 2019;3(1):e000006. doi:10.1136/bmjos-2018-000006

5. Bastings MM, Koudstaal S, Kieltyka RE, et al. A fast pH-switchable and self-healing supramolecular hydrogel carrier for guided, local catheter injection in the infarcted myocardium. Adv Heal Mater. 2014;3(1):70–78. doi:10.1002/adhm.201300076

6. Koudstaal S, Jansen of Lorkeers S, Gho JM, et al. Myocardial infarction and functional outcome assessment in pigs. J Vis Exp. 2014;(86). doi:10.3791/51269

7. Pape AC, Bakker MH, Tseng CC, et al. An Injectable and Drug-loaded Supramolecular Hydrogel for Local Catheter Injection into the Pig Heart. J Vis Exp. 2015;(100):e52450. doi:10.3791/52450

8. Feyen DAM, van den Akker F, Noort W, Chamuleau SAJ, Doevendans PA, Sluijter JPG. Isolation of Pig Bone Marrow-Derived Mesenchymal Stem Cells. In: Gnecchi M, ed. Mesenchymal Stem Cells: Methods and Protocols. New York, NY: Springer New York; 2016:225–232. doi:10.1007/978-1-4939-3584-0_12

9. Koudstaal S, Bastings MM, Feyen DA, et al. Sustained delivery of insulin-like growth factor-1/hepatocyte growth factor stimulates endogenous cardiac repair in the chronic infarcted pig heart. J Cardiovasc Transl Res. 2014;7(2):232–241. doi:10.1007/s12265-013-9518-4

10. Schotman MJG, Peters MMC, Krijger GC, et al. In Vivo Retention Quantification of Supramolecular Hydrogels Engineered for Cardiac Delivery. Adv Healthc Mater. 2021;10:2001987. doi:10.1002/adhm.202001987

11. van der Spoel TIG, Jansen of Lorkeers SJ, Agostoni P, et al. Human relevance of pre-clinical studies in stem cell therapy: systematic review and meta-analysis of large animal models of ischaemic heart disease. Cardiovasc Res. 2011;91(4):649–658. doi:10.1093/cvr/cvr113

12. Bakker MH, Tseng CCS, Keizer HM, et al. MRI Visualization of Injectable Ureidopyrimidinone Hydrogelators by Supramolecular Contrast Agent Labeling. Adv Healthc Mater. 2018;7(11):1701139. doi:https://doi.org/10.1002/adhm.201701139

13. Lyngbæk S, Ripa RS, Haack-Sørensen M, et al. Serial in vivo imaging of the porcine heart after human mesenchymal stromal cells. Int J Cardiovasc Imaging. 2010;26:273–284. doi:10.1007/s10554-009-9532-4

14. Collantes M, Pelacho B, García-Velloso MJ, et al. Non-invasive in vivo imaging of cardiac stem/progenitor cell biodistribution and retention after intracoronary and intramyocardial delivery in a swine model of chronic ischemia reperfusion injury. J Transl Med. 2017;15:56. doi:10.1186/s12967-017-1157-0

15. Hofmann M, Wollert KC, Meyer GP, et al. Monitoring of bone marrow cell homing into the infarcted human myocardium. Circulation. 2005;111(17):2198–2202. doi:10.1161/01.CIR.0000163546.27639.AA

16. Hudson W, Collins MC, deFreitas D, Sun YS, Muller-Borer B, Kypson AP. Beating and Arrested Intramyocardial Injections Are Associated with Significant Mechanical Loss: Implications for Cardiac Cell Transplantation. J Surg Res. 2007;142(2):263–267. doi:10.1016/j.jss.2007.03.021

17. Afzal MR, Samanta A, Shah ZI, et al. Adult bone marrow cell therapy for ischemic heart disease: Evidence and insights from randomized controlled trials. Circ Res. 2015;117(6):558–575. doi:10.1161/CIRCRESAHA.114.304792

18. Hou D, Youssef EA, Brinton TJ, et al. Radiolabeled Cell Distribution After Intramyocardial, Intracoronary, and Interstitial Retrograde Coronary. Circulation. 2005;112(9 suppl):I150-I156. doi:10.1161/CIRCULATIONAHA.104.526749

19. Perin EC, Tian M, Marini FC, et al. Imaging long-term fate of intramyocardially implanted mesenchymal stem cells in a porcine myocardial infarction model. PLoS One. 2011;6(9). doi:10.1371/journal.pone.0022949

